# E-Cigarette Synthetic Cooling Agent WS-23 and Nicotine Aerosols Differentially Modulate Airway Epithelial Cell Responses

**DOI:** 10.1101/2022.06.20.496868

**Authors:** Marko Manevski, Dinesh Devadoss, Shaiesh Yogeswaran, Irfan Rahman, Hitendra S. Chand

## Abstract

Electronic cigarette (e-cig) aerosol/vape exposures are strongly associated with pulmonary dysfunctions, and the airway epithelial cells (AECs) of respiratory passages play a pivotal role in understanding this association. However, not much is known about the effect of synthetic cooling agents such as WS-23 on AECs. WS-23 is a synthetic menthol-like cooling agent widely used to enhance the appeal of e-cigs and to suppress the harshness and bitterness of other e-cig constituents. Using primary human AECs, we compared the effects of aerosolized WS-23 with propylene glycol/vegetable glycerin (PG/VG) vehicle control and nicotine aerosol exposures. AECs treated with 3% WS-23 aerosols showed a significant increase in viable cell numbers compared to PG/VG-vehicle aerosol exposed cells and cell growth was comparable following 2.5% nicotine aerosol exposure. AEC inflammatory factors, IL-6 and ICAM-1 levels were significantly suppressed by WS-23 aerosols compared to PG/VG-controls. When differentiated AECs were challenged with WS-23 aerosols, there was a significant increase in MUC5AC+ goblet cells with no discernible change in SCGB1A1+ secretory cells. Compared to PG/VG-controls, WS-23 or nicotine aerosols presented with increased goblet cell numbers, but there was no synergistic effect of WS-23+nicotine combination exposure. Thus, WS-23 and nicotine aerosols modulate the AEC responses and induce goblet cell hyperplasia, which could impact the airway physiology and susceptibility to respiratory diseases.

## 1. INTRODUCTION

Electronic cigarettes (e-cigs) or other electronic nicotine delivery systems (ENDS) are marketed as products to aid tobacco smoking cessation. However, recent studies have found that e-cig/ENDS use during smoking cessation led to both a reduced effectiveness in cessation and a higher relapse rate as compared to no product use. (Kalkhoran and Glantz, 2016, Baenziger et al., 2021, Pierce et al., 2021, Wang et al., 2021, Chen et al., 2022) Menthol usage in e-cigarettes, e-liquids, and other ENDS products is known to enhance the appeal of these products, particularly to young adult users, as it reduces the harshness and bitterness of the products. (Gaiha et al., 2022, FDA, 2021) The U.S. Food and Drug Administration (FDA) recently banned flavorings other than menthol and tobacco in closed pod systems, dramatically increasing the use of menthol-containing e-cig/ENDS products. (FDA, 2020, Wang et al., 2020b) Reports have suggested that this change may have led to increased presence of menthol and synthetic cooling agents such as WS-23 in e-cig products which may ultimately exposure users to more harm. (Omaiye et al., 2021) Chemical analysis of various menthol-flavored e-cig products corroborated the presence of various harmful compounds (Omaiye et al., 2021, Jabba and Jordt, 2019, Gerloff et al., 2017, Kaur et al., 2020). Despite efforts to curb the appeal of e-cigarettes, particularly to young adults, the use of menthol and “iced/cooling” flavors has only increased in popularity and has potentially contributed to the increased addictive properties of e-cigarettes. (Kaur et al., 2020) Considering the recent outbreak of e-cigarette or vaping use-associated lung injury (EVALI) (Chand et al., 2019), the current rate of e-cig use may cause severe comorbid conditions among a larger population. Among various cooling agents analyzed, the synthetic cooling agent WS-23 was reportedly used most prominently. (Jabba et al., 2022) 2-isopropyl-N,2,3-trimethylbutyramide, commonly known as WS-23, was found to be present in most e-liquids marketed in the U.S. in quantities that may exceed consumer exposure safety standards. (Jabba et al., 2022)

Although studies on the biological effects of WS-23 are lacking, a recent report found that there may be cytotoxicity induced by WS-23 exposure in vitro. (Omaiye et al., 2021) One study found that various flavoring products induced ROS generation and superoxide production in vitro in lung epithelial cell lines and monocytes. (Muthumalage et al., 2019) Similarly, ROS generation and pro-inflammatory effects were observed to be induced by e-cig aerosol/vape exposure; these effects were further amplified by flavored e-cigs in periodontal fibroblasts. (Sundar et al., 2016) Furthermore, e-cig use affected lung inflammatory responses, and importantly, the aerosols consisting of propylene glycol/vegetable glycerin (PG/VG) vehicle alone were found to induce a potent pro-inflammatory response and immune infiltration in bronchoalveolar lavage fluid (BALF). (Wang et al., 2020a)

The respiratory airway epithelial cells (AECs) are pivotal to innate immune defense against inhaled toxicants/allergens, and the AEC responses to aerosolized e-cig components are crucial for orchestrating the lung immune responses. (Devadoss et al., 2019, Manevski et al., 2020b) Any dysregulation in AEC-mediated responses can significantly impact the susceptibility to infection (Devadoss et al., 2021a); a recent study has shown that e-cig use induces a reduction in AEC ciliary beating frequency, as well as changes in cytokine and chemokine production. (Jasper et al., 2021) Nevertheless, the impact of synthetic cooling agents such as WS-23 individually has not been investigated.

With the continuously increasing usage of the WS-23 cooling agent, coupled with the increased usage of e-cigarettes, particularly in young adults, there is a need to assess the effects of WS-23 aerosols/vapes on both AECs and the innate immune response of airways.

In this report, we analyzed the effect of aerosolized WS-23 in vitro using primary airway epithelial cells and compared the responses to those induced by PG/VG vehicle aerosols using the Buxco EVT exposure system (Data Sciences International, St. Paul, MN). Notably, the PG/VG vehicle has been demonstrated to have an aberrant effect on lipid homeostasis and downregulate innate immune responses in AECs, particularly against viral infection. (Madison et al., 2019) As such, we compared the effect of WS-23 against PG/VG vehicle and found that WS-23 may be reducing the expression of interleukin (IL)-6 and ICAM-1, thereby dysregulating the AEC innate immune responses.

## 2. MATERIALS and METHODS

### 2.1 Human Airway Epithelial Cell Culture

Primary human AECs were seeded in clear TC-treated 6-well plates (Corning Costar^®^) using bronchial epithelial cell growth media (BEGM, Lonza, or UNC MLI Cell Culture Core), and e-cig aerosol treatments were started 24 hours after seeding. Alternatively, primary AECs were also grown in air-liquid interface (ALI) cultures as described previously (Devadoss et al., 2021b) and cells were differentiated for a minimum of 21 days before treatments.

### 2.2 E-liquid Reagents and E-cig Aerosol/Vape Exposures

E-liquid synthetic cooling and flavoring agent WS-23 (CAS#51115-67-4, from FlavorJungle, Bellingham, WA) was used with or without Nicotine (Sigma-Aldrich, Inc.), and PG/VG (1:1, propylene glycol: vegetable glycerin) was used as a vehicle control. There were four treatment groups, where cells were treated with PG/VG, 3% WS-23 in PG/VG, 2.5% Nicotine in PG/VG, or 2.5% Nicotine + 3% WS-23 prepared in PG/VG. Human AECs were exposed to e-liquid aerosols using the Buxco E-cigarette, Vapor, and Tobacco (EVT) exposure system (Data Sciences International, St. Paul, MN, USA) as described before (Yogeswaran and Rahman, 2022). Briefly, cells were exposed to e-cig aerosols for 15 minutes per day with a puff topography of 2 puffs/minute (3 s/puff, 55 mL/puff). Smok^®^ X-Priv mod kit was used for smoke delivery installed with Prince^®^ V12 triple mesh coils with 90 watts coil wattage. After exposure, cells were incubated at 37°C and 5% CO_2_ for another 24 hours and cells and media supernatant were harvested after 48 and 72 hours of exposure. Cell viability was assessed by the trypan blue exclusion method. Briefly, trypsinized cells were resuspended in phosphate-buffered saline (PBS), and samples were mixed 1:1 with 0.4% trypan blue solution (catalog no. 302643; Sigma-Aldrich), and live/dead cells were counted using TC20 automated cell counter (Bio-Rad Inc).

### 2.3 Inflammatory Gene Expression Analysis by qRT-PCR

Total RNA extraction was performed using the RNeasy Mini kit (Qiagen) according to the manufacturer’s instructions, and cDNAs were synthesized using the Applied Biosciences High-Capacity RNA-to-cDNA^®^ Synthesis Kit (Thermo Fisher Scientific, Inc), per manufacturer’s instructions. Expression levels of *ICAM-1* and *IL6* mRNAs were quantified using SYBR Green-based primers and the iTaq master mix (Bio-Rad Inc) in the Bio-Rad CFX Real-Time PCR detection system (Bio-Rad Inc). Relative quantification data were obtained using the delta-delta (ΔΔ)Ct method by normalizing to the respective *beta-actin* and/or *GAPDH* mRNA levels as described recently. (Devadoss et al., 2021b)

### 2.4 Secretory Inflammatory Factor Analysis by ELISA

The protein levels of ICAM-1 and IL-6 were determined using human ELISA kits against ICAM-1 (LifeSpan Biosciences Inc., Seattle, WA) and IL-6 (BioLegend Inc., San Diego, CA), respectively, as per manufacturers’ instructions.

### 2.5 Immunocytochemical staining and Imaging Analysis

For immunocytochemical staining, cells were fixed with 4% paraformaldehyde (PFA) and washed in 0.05% v Brij-35 in PBS (pH 7.4) and immunostained using antibodies to MUC5AC (Millipore Inc., Burlington, MA), SCGB1A1 or secretoglobulin 1A1 (Santa Cruz Biotechnology, Santa Cruz, CA) and β-tubulin (Cell Signaling Tech., Danvers, MA) or isotype controls. Briefly, cells were blocked using a solution containing 3% BSA, 1% Gelatin, and 1% normal donkey serum with 0.1% Triton X-100 and 0.1% Saponin and were stained with antibodies. The immunolabelled cells were detected using respective secondary antibodies conjugated fluorescent dyes (Jackson ImmunoResearch Lab Inc., West Grove, PA) and mounted with 4’,6-diamidino-2-phenylindole (DAPI) containing Fluormount-G^TM^ (SouthernBiotech, Birmingham, AL) for nuclear staining. Immunofluorescent images were captured using the BZX700 Microscopy system (Keyence Corp., Japan) and analyzed using NIH Image J software.

### 2.6 Statistical Analysis

Data expressed as mean±SEM were analyzed using GraphPad Prism Software (GraphPad Software Inc.) using one-way analysis of variance (ANOVA) with and following Tukey’s multiple comparison test. When significant main effects were detected (*p*<0.05), student’s t-test was used to determine differences between the groups. Studies were performed following three separate experiments.

## 3. RESULTS & DISCUSSION

### 3.1 Synthetic cooling agent WS-23 aerosols modulate human airway epithelial cell viability

We first analyzed the effects of aerosolized 3% WS-23 on cell viability of AECs in submerged cultures at 48 and 72 h of treatment. The viable cell counts showed no significant change in cell numbers among all the groups tested. However, the WS-23 aerosol-treated cells showed a trend toward reducing viable cell numbers (**Figure 1A**). Interestingly, at 72 h of treatment, specifically in cells treated with 3% WS-23 or 2.5% nicotine aerosols, we observed a significant increase in viable cell numbers, but there was no synergistic effect observed in cells treated with WS-23+nicotine combination (**Figure 1B**). It is noteworthy that cells treated with PG/VG aerosols showed a marked reduction in cell numbers at 72 h compared to 48 h of treatment.

**Figure 1.**
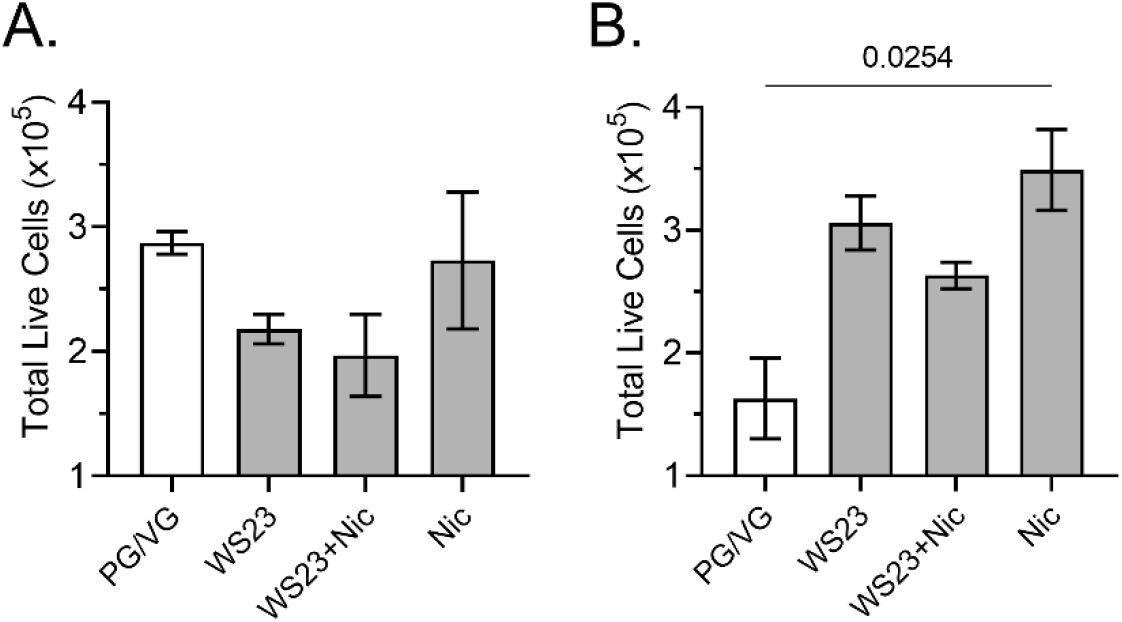
Synthetic cooling agent WS-23 aerosols induce cell proliferation in AECs following 72 h of treatment. Live cell numbers following (**A.**) 48h, and (**B.**) 72h treatment with aerosolized PG/VG (50:50), PG/VG + 3% WS-23, PG/VG + 3% WS-23 + 2.5% nicotine, or PG/VG + 2.5% nicotine using Buxco EVT system. Data shown as mean ±SEM and analyzed by one-way ANOVA, n=2/gp.

### 3.2 PG/VG vehicle induces an inflammatory response in human AECs independent of nicotine and WS-23

As PG/VG has been shown to dysregulate AEC responses (Madison et al., 2019) we analyzed *IL-6* and *ICAM-1* mRNA levels upon exposure to PG/VG for 24h, 48h, and 72h.Interestingly, we observed a significant increase in both *IL-6* and *ICAM-1* mRNA at 48h of exposure, which was further potentiated at 72h of exposure time (**Figure 2A–2B**). These data suggest that PG/VG vehicle without added constituents can dysregulate the cellular responses, which may be further potentiated or inhibited by the addition of nicotine or WS-23.

**Figure 2.**
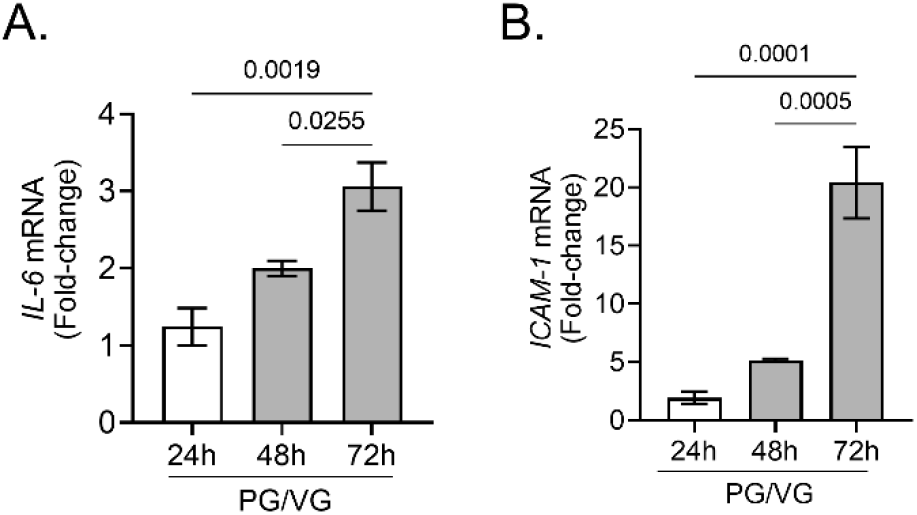
E-liquid vehicle (PG/VG) aerosols induce *IL-6* and *ICAM-1* mRNA expression in AECs. (**A.**) *IL-6* and (**B.**) *ICAM-1* mRNA levels following 24h, 48h, and 72h of exposure to PG/VG vehicle aerosols. Data shown as mean±SEM and analyzed by one-way ANOVA, n=2/gp.

### 3.3 WS-23 aerosols alter innate inflammatory response kinetics of human AECs

We next analyzed the effects of WS-23 aerosols on AEC mRNA expression of inflammatory factors, *IL-6,* and *ICAM-1,* which are important modulators of AEC innate responses (Devadoss et al., 2021b). At 48 h of treatment, *IL-6* mRNA levels were significantly reduced by WS-23 or nicotine aerosol exposure, as the WS-23, WS-23+nicotine, and nicotine alone aerosol treated groups presented with 2.0-fold or higher reduction in *IL-6* mRNA levels, compared to PG/VG-treated controls (**Figure 3A**). In contrast, cells treated with WS-23 or nicotine aerosols showed a trend towards increased *ICAM-1* mRNA expression; however, WS-23+nicotine combined treatment induced a significant increase in *ICAM-1* expression (**Figure 3B**). Thus, *ICAM-1* mRNA levels were increased by WS-23 and nicotine treatments, and treatment with both WS-23 and nicotine in conjunction (WS-23+nicotine) further potentiated the increase in *ICAM-1* mRNA levels.

**Figure 3.**
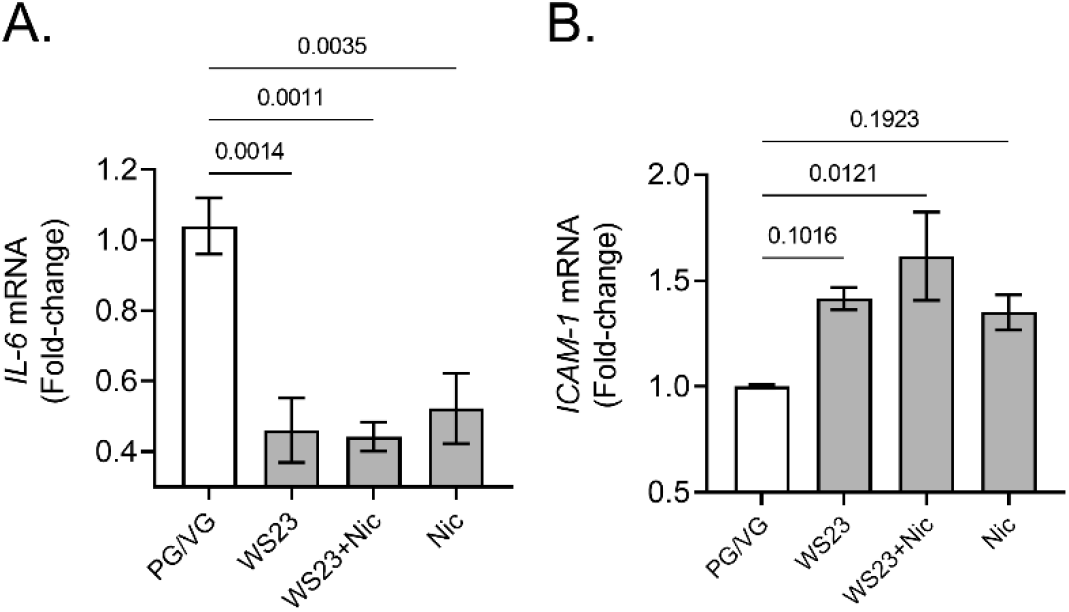
WS-23 e-liquid aerosols suppress the expression of *IL-6* and induce the expression of *ICAM-1* mRNA expression following 48 h of treatment. Relative quantities of (**A.**) *IL-6* and (**B.**) *ICAM-1* mRNAs following 48 h exposure to aerosolized PG/VG, PG/VG + 3% WS-23, PG/VG + 3% WS-23 + 2.5% nicotine, or PG/VG + 2.5% nicotine. Data shown as mean±SEM and analyzed by one-way ANOVA, n=2/gp.

After 72 h treatment, we observed a similar trend in the expression levels of *IL-6* mRNA. Cells treated with WS-23, WS-23+nicotine, and nicotine alone aerosols presented with a 1.5-, 1.7-, and 2.0-fold decrease in *IL-6* mRNA levels, respectively, compared to PG/VG-treated controls after 72 h of treatment (**Figure 4A**). Surprisingly, *ICAM-1* mRNA levels were markedly reduced, with WS-23, WS-23 + nicotine, and nicotine alone treatments causing a 1.8-, 1.75-, and 1.88-fold reduction in *ICAM-1* expression, respectively, when compared to the PG/VG-treated controls (**Figure 4B**).

**Figure 4.**
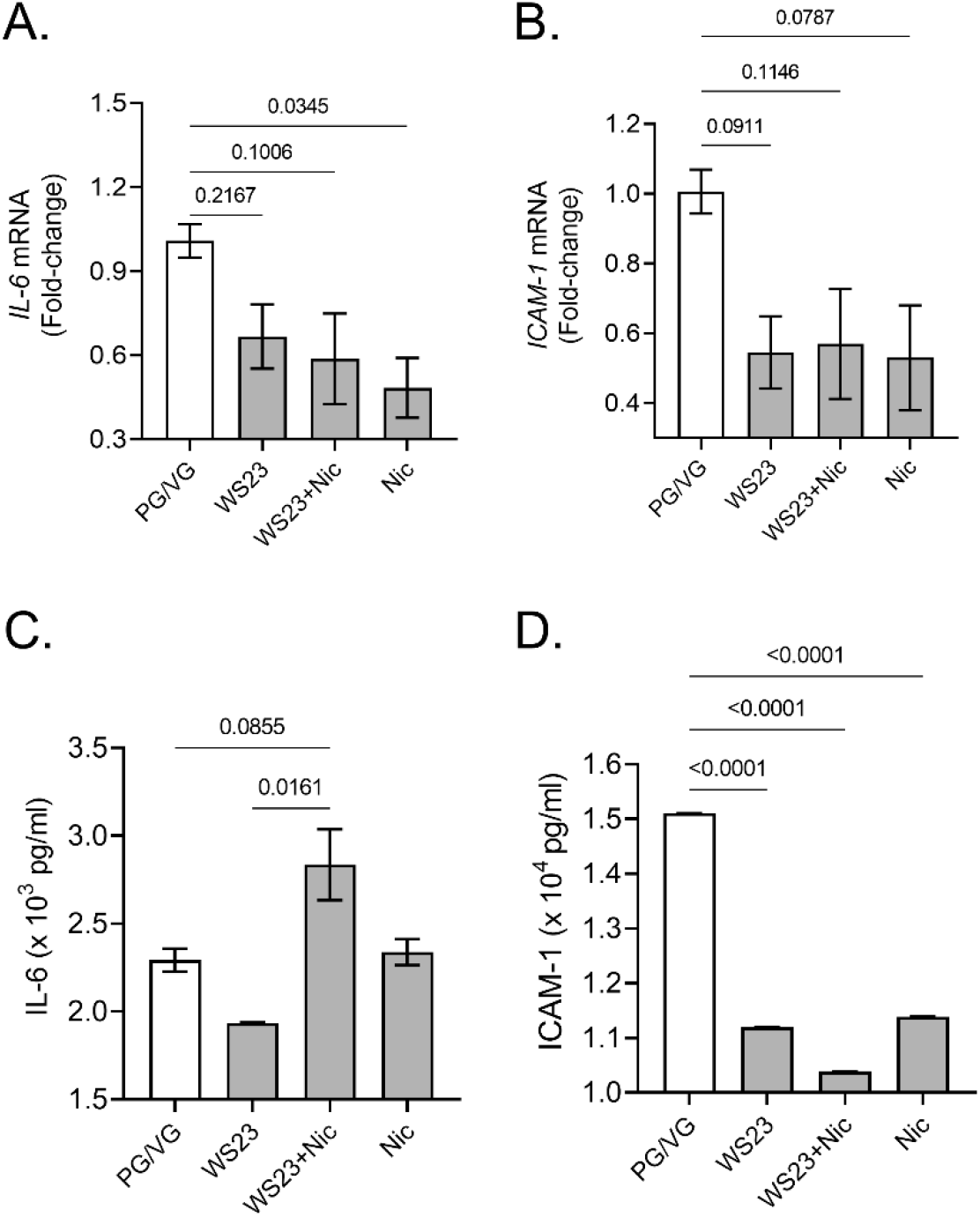
Aerosolized WS-23 exposure suppresses IL-6 and ICAM-1 expression and secretion in conjunction with nicotine aerosol exposure. Relative quantity of (**A.**) *IL-6,* and (**B.**) *ICAM-1* mRNA levels following 72h of exposure as evaluated by qRT-PCR. Secreted protein levels of (**C.**) IL-6, and (**D.**) ICAM-1 as evaluated in cell culture supernatants by specific ELISA assays after 72h of aerosol exposure. Data shown as mean±SEM and analyzed by one-way ANOVA, n=2/gp.

We next corroborated these results by investigating the changes in protein levels of secretory IL-6 and ICAM-1 in culture media supernatants harvested at 72 h treatment. Interestingly, IL-6 secretory levels were on an average 2,292 pg/ml in PG/VG control culture media; and 1,932 pg/ml in WS-23; 2,835 pg/ml in WS-23+nicotine; and 2,338 pg/ml in nicotine treated groups (**Figure 4C**). There was a significant reduction in ICAM-1 expression upon all three treatments. The PG/VG control treatment presented with 15,105 pg/ml of ICAM-1, whereas the WS-23, WS-23+nicotine, and nicotine alone treatments presented an average of 11,196, 10,381, and 11,388 pg/ml, respectively (**Figure 4D**). Thus, aerosolized synthetic cooling agent WS-23 alters the innate airway immune responses of human AECs.

### 3.4 WS-23 aerosol exposure modulates the goblet cell differentiation in AECs

Next, we analyzed the effects of WS-23 aerosols on a differentiated AEC population cultured on an air-liquid interface. Groups of Transwells were treated with aerosolized PG/VG vehicle, WS-23, nicotine, and WS23+nicotine. After 72 h treatment, there were significant changes in MUC5AC+ mucous/goblet cells by both WS-23 or nicotine aerosol exposure (**Figure 5A**). The WS-23 and nicotine alone aerosol treated groups presented with increased goblet cells compared to PG/VG-treated controls, but there was no synergistic effect of WS-23+nicotine combination exposure (**Figure 5B**). In contrast, there was a reduction in SCGB1A1 + secretory club cells in WS-23, WS-23+nicotine, or nicotine aerosols treated groups (**Figures 5C** and **5D**). Thus, WS-23 and nicotine aerosols modulate the airway secretory cell population by affecting the goblet cell hyperplasia, which could impact the respiratory physiology and needs further validation using in vivo exposure models.

**Figure 5.**
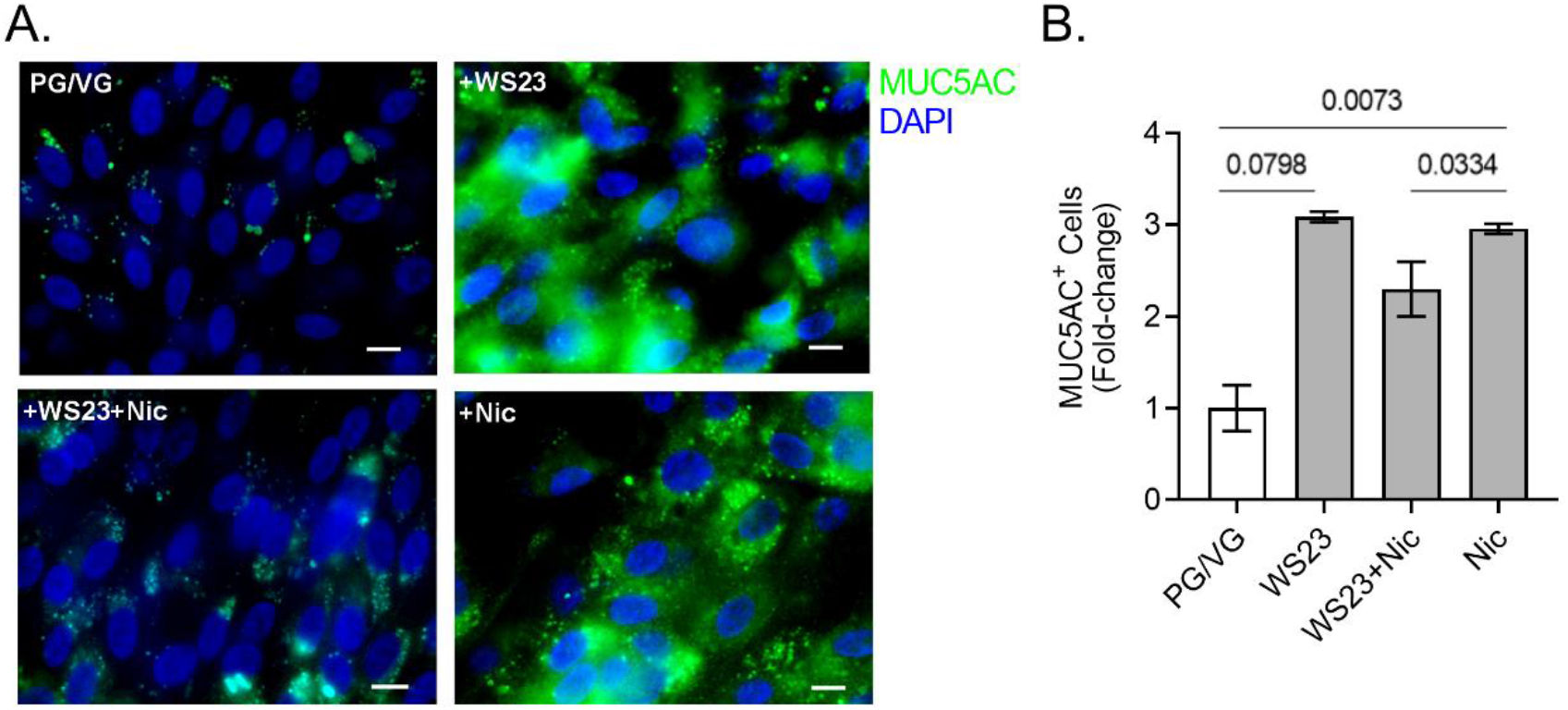
Aerosolized WS-23 exposure augments MUC5AC+ goblet cell hyperplasia in air-liquid interface differentiated AECs. (**A.**) Representative micrographs showing MUC5AC immunopositivity (shown in green) in AECs treated with aerosolized PG/VG, PG/VG + 3% WS-23, PG/VG + 3% WS-23 + 2.5% nicotine, or PG/VG + 2.5% nicotine; and nuclei were stained with DAPI (shown in blue), scale–5μm. **(B.)** Quantification of MUC5AC-positive (MUC5AC^+^) goblet cells in each treatment group. Data shown as mean±SEM and analyzed by one-way ANOVA, n=2/gp.

## 4. DISCUSSION

The FDA considers flavoring and cooling agents safe when utilized as food additives; however, the risks associated with their inhalation, through vaping, are poorly defined. (Hallagan, 2015) Little is known about how e-cig constituents affect the respiratory tract, specifically when emerging evidence indicates that the acute effects of e-cig products use on the respiratory system need to be revisited (Viswam et al., 2018, Khan et al., 2018, Sommerfeld et al., 2018, Layden et al., 2019, Madison et al., 2019). Furthermore, it has been reported that menthol, in concentrations found in e-cig aerosols may disturb cell homeostasis and can trigger oxidative stress via the NF-κB pathway. (Nair et al., 2020) Induction of oxidative stress and chronic mitochondrial dysregulation is central in many pathologic conditions such as chronic inflammatory and aging-associated degenerative diseases. (Manevski et al., 2020a) Most importantly, alarmingly high toxic levels of synthetic cooling agents/coolants such as WS-3 or WS-23 carboxamides are used in the emerging e-cigs products. Still, the risk associated with their inhalation and safety regulations is understudied. These are not only found in mint/menthol-flavored products but also in the fruit-, candy-, and ice flavors, including the popular disposable pod-and mod-based products. Even without nicotine or flavoring agents, e-cig use can induce significant changes in the lung epithelium; one study found that chronic exposure to e-cig aerosols in a mouse model suppresses the innate immune response, particularly against viral infection. (Madison et al., 2019) Aerosolized e-liquid solvents exposure induces significant changes in the airway epithelium, with or without added flavoring elements. However, the results are highly variable. Contrasting changes in inflammatory factors expression were reported on a significant increase either in expression or with no change in the expression levels of IL-6 and CXCL-8 (Jasper et al. 2021). This discrepancy could be mainly attributed to the exposure systems used for e-cig aerosols and the addition of nicotine or other flavoring chemicals. (Jasper et al., 2021) There have been significant variations in study design, and the need for further investigation is dire. As newer ENDS emerge and regulations continue to change with evolving consumer preferences, it is crucial to understand the effects of prominently used flavoring compounds, like WS-23. Our results further indicate that PG/VG dysregulates cellular responses with or without nicotine or flavoring agents. Notably, we found that those responses were inhibited by adding additional constituents such as WS-23 and nicotine.

In this study, we used a submerged culture model of primary AECs and exposed them to e-cig vapor using PG/VG as a vehicle-only control and compared the effects to those with PG/VG and WS-23, nicotine or WS-23 and nicotine exposure. Our results demonstrated that even at 48 hours after exposure, there is a significant reduction in the expression levels of IL-6, and concurrently, a trend of increased expression of ICAM-1, with a significant increase in ICAM-1 levels upon WS-23+nicotine treatment. These data further corroborate the observations that WS-23 may alter the innate immune responses of AEC. Suppressing the IL-6 levels may hinder the rapid AEC immune response to pathogen presence. (Jones and Jenkins, 2018) Alternatively, the increased levels of ICAM-1 suggest a differential effect on regulatory pathways. It has been reported that e-cig exposure without nicotine may induce a transient increase in secretory ICAM-1 levels; however, the effects of WS-23 on these pathways have not been thoroughly investigated. (Chatterjee et al., 2019) Furthermore, at the 72-hour time point, we observed a significant reduction in the expression of *IL-6* mRNA levels, the expression of *ICAM-1* mRNA levels, and the secretory protein levels of ICAM-1. Research evaluating the effect of WS-23 on MUC5AC+ goblet cells and airway epithelial cells is lacking; however, it has been shown that WS-23 may induce MUC5AC expression in patients with chronic obstructive pulmonary disease (COPD) by binding to transient receptor potential cation channel (TRMP)8, which is upregulated in COPD patients. (Li et al., 2005, Kaneko and Szallasi, 2014) Nicotine exposure has been well established as an inducer of mucin MUC5AC expression. (Escobar et al., 2021, Haswell et al., 2021) Here, we report that WS-23 may be inducing a similar response independent of nicotine presence and may require further investigation into long-term effects.

Interestingly, it has been reported that nicotine can reduce the levels of SCGB1A1 or CCSP by reducing the transcription factor FOXA2. (Warren and O’ Reilly, 2019, Zhu et al., 2019) However, research investigating the effect of WS-23 on SCGB1A1 expression is lacking. We found that exposure to both WS-23 and nicotine induced a trend of increased SCGB1A1 expression compared to PG/VG control; however, these results were not significant (data not shown). This may suggest that WS-23 induces an acute increase in secretory cells, and further investigation is required into the long-term changes that may be induced by WS-23 exposure, independent of nicotine.

Our data further corroborate the effects of both nicotine and WS-23 on AECs and suggest that flavoring agents may amplify or reduce the toxic effects of various e-cig components such as those induced by PG/VG vehicle and nicotine or nicotine salt additives. Notably, WS-23 and nicotine presence reduced the inflammatory responses that were strongly induced by PG/VG vehicle independently. These data collectively suggest that aerosolized synthetic cooling agent WS-23 alters the innate airway immune responses of human AECs and thus, potentially could increase the susceptibility to respiratory pathologies.

## AUTHOR CONTRIBUTIONS

MM and DD designed and conducted the experiments; MM and HSC wrote, edited, and revised the manuscript; MM and DD were responsible for data curation; IR provided the oversight on study design and edited the manuscript. SY provided technical suggestions and edited the manuscript. HSC and IR conceptually designed the study and manuscript and acquired funding.

## INSTITUTIONAL REVIEW BOARD STATEMENT

The procurement of human airway epithelial cells was approved by the Materials Transfer Agreement and Procurement. Exposure studies were performed under the laboratory protocols approved by Institutional Biosafety Committee (IBC) at the Florida International University.

## ACKNOWLEDGEMENTS

The authors would like to acknowledge Ariane Chung and Francisco De Leon for their assistance in the laboratory experiments.

## REFERENCES

1. Baenziger, O. N., Ford, L., Yazidjoglou, A., Joshy, G. & Banks, E. 2021. E-cigarette use and combustible tobacco cigarette smoking uptake among non-smokers, including relapse in former smokers: umbrella review, systematic review and meta-analysis. BMJ Open, 11, e045603. 10.1136/bmjopen-2020-045603.

2. Chand, H. S., Muthumalage, T., Maziak, W. & Rahman, I. 2019. Pulmonary Toxicity and the Pathophysiology of Electronic Cigarette, or Vaping Product, Use Associated Lung Injury. Front Pharmacol, 10, 1619. 10.3389/fphar.2019.01619.

3. Chatterjee, S., Tao, J.-Q., Johncola, A., Guo, W., Caporale, A., Langham, M. C. & Wehrli, F. W. 2019. Acute exposure to e-cigarettes causes inflammation and pulmonary endothelial oxidative stress in nonsmoking, healthy young subjects. American journal of physiology. Lung cellular and molecular physiology, 317, L155–L166. 10.1152/ajplung.00110.2019.

4. Chen, R., Pierce, J. P., Leas, E. C., Benmarhnia, T., Strong, D. R., White, M. M., Stone, M., Trinidad, D. R., Mcmenamin, S. B. & Messer, K. 2022. Effectiveness of e-cigarettes as aids for smoking cessation: evidence from the PATH Study cohort, 2017-2019. Tob Control. 10.1136/tobaccocontrol-2021-056901.

5. Devadoss, D., Acharya, A., Manevski, M., Pandey, K., Borchert, G. M., Nair, M., Mirsaeidi, M., Byrareddy, S. N. & Chand, H. S. 2021a. Distinct Mucoinflammatory Phenotype and the Immunomodulatory Long Noncoding Transcripts Associated with SARS-CoV-2 Airway Infection. medRxiv. 10.1101/2021.05.13.21257152.

6. Devadoss, D., Daly, G., Manevski, M., Houserova, D., Hussain, S. S., Baumlin, N., Salathe, M., Borchert, G. M., Langley, R. J. & Chand, H. S. 2021b. A long noncoding RNA antisense to ICAM-1 is involved in allergic asthma associated hyperreactive response of airway epithelial cells. Mucosal Immunol, 14, 630–639. 10.1038/s41385-020-00352-9.

7. Devadoss, D., Long, C., Langley, R. J., Manevski, M., Nair, M., Campos, M. A., Borchert, G., Rahman, I. & Chand, H. S. 2019. Long Noncoding Transcriptome in Chronic Obstructive Pulmonary Disease. Am J Respir Cell Mol Biol, 61, 678–688. 10.1165/rcmb.2019-0184TR.

8. Escobar, Y. H., Morrison, C. B., Chen, Y., Hickman, E., Love, C. A., Rebuli, M. E., Surratt, J. D., Ehre, C. & Jaspers, I. 2021. Differential responses to e-cig generated aerosols from humectants and different forms of nicotine in epithelial cells from nonsmokers and smokers. Am J Physiol Lung Cell Mol Physiol, 320, L1064–l1073. 10.1152/ajplung.00525.2020.

9. FDA 2020. FDA fnalizes enforcement policy on unauthorized favored cartridge-based e-cigarettes that appeal to children, including fruit and mint. FDA.

10. FDA 2021. Menthol and Other Flavors in Tobacco Products..FDA.

11. Gaiha, S. M., Lempert, L. K., Mckelvey, K. & Halpern-Felsher, B. 2022. E-cigarette devices, brands, and flavors attract youth: Informing FDA’s policies and priorities to close critical gaps. Addict Behav, 126, 107179. 10.1016/j.addbeh.2021.107179.

12. Gerloff, J., Sundar, I. K., Freter, R., Sekera, E. R., Friedman, A. E., Robinson, R., Pagano, T. & Rahman, I. 2017. Inflammatory Response and Barrier Dysfunction by Different e-Cigarette Flavoring Chemicals Identified by Gas Chromatography-Mass Spectrometry in e-Liquids and e-Vapors on Human Lung Epithelial Cells and Fibroblasts. Appl In Vitro Toxicol, 3, 28–40. 10.1089/aivt.2016.0030.

13. Hallagan, J. 2015. The Safety Assessment and Regulatory Authority to Use Flavors - Focus on E-Cigarettes. Available: https://www.femaflavor.org/node/24344 [Accessed 18 May 2021].

14. Haswell, L. E., Smart, D., Jaunky, T., Baxter, A., Santopietro, S., Meredith, S., Camacho, O. M., Breheny, D., Thorne, D. & Gaca, M. D. 2021. The development of an in vitro 3D model of goblet cell hyperplasia using MUC5AC expression and repeated whole aerosol exposures. Toxicol Lett, 347, 45–57. 10.1016/j.toxlet.2021.04.012.

15. Jabba, S. V., Erythropel, H. C., Torres, D. G., Delgado, L. A., Woodrow, J. G., Anastas, P. T., Zimmerman, J. B. & Jordt, S. E. 2022. Synthetic Cooling Agents in US-marketed E-cigarette Refill Liquids and Popular Disposable Ecigarettes: Chemical Analysis and Risk Assessment. Nicotine Tob Res. 10.1093/ntr/ntac046.

16. Jabba, S. V. & Jordt, S. E. 2019. Risk Analysis for the Carcinogen Pulegone in Mint-and Menthol-Flavored e-Cigarettes and Smokeless Tobacco Products. JAMA Intern Med, 179, 1721–1723. 10.1001/jamainternmed.2019.3649.

17. Jasper, A. E., Sapey, E., Thickett, D. R. & Scott, A. 2021. Understanding potential mechanisms of harm: the drivers of electronic cigarette-induced changes in alveolar macrophages, neutrophils, and lung epithelial cells. Am J Physiol Lung Cell Mol Physiol, 321, L336–l348. 10.1152/ajplung.00081.2021.

18. Jones, S. A. & Jenkins, B. J. 2018. Recent insights into targeting the IL-6 cytokine family in inflammatory diseases and cancer. Nat Rev Immunol, 18, 773–789. 10.1038/s41577-018-0066-7.

19. Kalkhoran, S. & Glantz, S. A. 2016. E-cigarettes and smoking cessation in real-world and clinical settings: a systematic review and meta-analysis. Lancet Respir Med, 4, 116–28. 10.1016/s2213-2600(15)00521-4.

20. Kaneko, Y. & Szallasi, A. 2014. Transient receptor potential (TRP) channels: a clinical perspective. Br J Pharmacol, 171, 2474–507. 10.1111/bph.12414.

21. Kaur, G., Gaurav, A., Lamb, T., Perkins, M., Muthumalage, T. & Rahman, I. 2020. Current Perspectives on Characteristics, Compositions, and Toxicological Effects of E-Cigarettes Containing Tobacco and Menthol/Mint Flavors. Front Physiol, 11, 613948. 10.3389/fphys.2020.613948.

22. Khan, M. S., Khateeb, F., Akhtar, J., Khan, Z., Lal, A., Kholodovych, V. & Hammersley, J. 2018. Organizing pneumonia related to electronic cigarette use: A case report and review of literature. Clin Respir J, 12, 1295–1299. 10.1111/crj.12775.

23. Layden, J. E., Ghinai, I., Pray, I., Kimball, A., Layer, M., Tenforde, M., Navon, L., Hoots, B., Salvatore, P. P., Elderbrook, M., Haupt, T., Kanne, J., Patel, M. T., Saathoff-Huber, L., King, B. A., Schier, J. G., Mikosz, C. A. & Meiman, J. 2019. Pulmonary Illness Related to E-Cigarette Use in Illinois and Wisconsin - Preliminary Report. N Engl J Med. 10.1056/NEJMoa1911614.

24. Li, Y., Jia, Y. C., Cui, K., Li, N., Zheng, Z. Y., Wang, Y. Z. & Yuan, X. B. 2005. Essential role of TRPC channels in the guidance of nerve growth cones by brain-derived neurotrophic factor. Nature, 434, 894–8. 10.1038/nature03477.

25. Madison, M. C., Landers, C. T., Gu, B. H., Chang, C. Y., Tung, H. Y., You, R., Hong, M. J., Baghaei, N., Song, L. Z., Porter, P., Putluri, N., Salas, R., Gilbert, B. E., Levental, I., Campen, M. J., Corry, D. B. & Kheradmand, F. 2019. Electronic cigarettes disrupt lung lipid homeostasis and innate immunity independent of nicotine. J Clin Invest, 129, 4290–4304. 10.1172/jci128531.

26. Manevski, M., Devadoss, D., Castro, R., Delatorre, L., Yndart, A., Jayant, R. D., Nair, M. & Chand, H. S. 2020a. Development and Challenges of Nanotherapeutic Formulations for Targeting Mitochondrial Cell Death Pathways in Lung and Brain Degenerative Diseases. Crit Rev Biomed Eng, 48, 137–152. 10.1615/CritRevBiomedEng.2020034546.

27. Manevski, M., Muthumalage, T., Devadoss, D., Sundar, I. K., Wang, Q., Singh, K. P., Unwalla, H. J., Chand, H. S. & Rahman, I. 2020b. Cellular stress responses and dysfunctional Mitochondrial-cellular senescence, and therapeutics in chronic respiratory diseases. Redox Biol, 33, 101443. 10.1016/j.redox.2020.101443.

28. Muthumalage, T., Lamb, T., Friedman, M. R. & Rahman, I. 2019. E-cigarette flavored pods induce inflammation, epithelial barrier dysfunction, and DNA damage in lung epithelial cells and monocytes. Sci Rep, 9, 19035. 10.1038/s41598-019-51643-6.

29. Nair, V., Tran, M., Behar, R. Z., Zhai, S., Cui, X., Phandthong, R., Wang, Y., Pan, S., Luo, W., Pankow, J. F., Volz, D. C. & Talbot, P. 2020. Menthol in electronic cigarettes: A contributor to respiratory disease? Toxicol Appl Pharmacol, 407, 115238. 10.1016/j.taap.2020.115238.

30. Omaiye, E. E., Luo, W., Mcwhirter, K. J., Pankow, J. F. & Talbot, P. 2021. Flavour chemicals, synthetic coolants and pulegone in popular mint-flavoured and menthol-flavoured e-cigarettes. Tob Control. 10.1136/tobaccocontrol-2021-056582.

31. Pierce, J. P., Chen, R., Kealey, S., Leas, E. C., White, M. M., Stone, M. D., Mcmenamin, S. B., Trinidad, D. R., Strong, D. R., Benmarhnia, T. & Messer, K. 2021. Incidence of Cigarette Smoking Relapse Among Individuals Who Switched to e-Cigarettes or Other Tobacco Products. JAMA Netw Open, 4, e2128810. 10.1001/jamanetworkopen.2021.28810.

32. Sommerfeld, C. G., Weiner, D. J., Nowalk, A. & Larkin, A. 2018. Hypersensitivity Pneumonitis and Acute Respiratory Distress Syndrome From E-Cigarette Use. Pediatrics, 141. 10.1542/peds.2016-3927.

33. Sundar, I. K., Javed, F., Romanos, G. E. & Rahman, I. 2016. E-cigarettes and flavorings induce inflammatory and pro-senescence responses in oral epithelial cells and periodontal fibroblasts. Oncotarget, 7, 77196–77204. 10.18632/oncotarget.12857.

34. Viswam, D., Trotter, S., Burge, P. S. & Walters, G. I. 2018. Respiratory failure caused by lipoid pneumonia from vaping e-cigarettes. BMJ Case Rep, 2018. 10.1136/bcr-2018-224350.

35. Wang, Q., Sundar, I. K., Li, D., Lucas, J. H., Muthumalage, T., Mcdonough, S. R. & Rahman, I. 2020a. E-cigarette-induced pulmonary inflammation and dysregulated repair are mediated by nAChR α7 receptor: role of nAChR α7 in SARS-CoV-2 Covid-19 ACE2 receptor regulation. Respir Res, 21, 154. 10.1186/s12931-020-01396-y.

36. Wang, R. J., Bhadriraju, S. & Glantz, S. A. 2021. E-Cigarette Use and Adult Cigarette Smoking Cessation: A Meta-Analysis. Am J Public Health, 111, 230–246. 10.2105/ajph.2020.305999.

37. Wang, T. W., Neff, L. J., Park-Lee, E., Ren, C., Cullen, K. A. & King, B. A. 2020b. Ecigarette Use Among Middle and High School Students - United States, 2020. MMWR Morb Mortal Wkly Rep, 69, 1310–1312. 10.15585/mmwr.mm6937e1.

38. Warren, R. & O’Reilly, M. A. 2019. An Elusive Fox that Suppresses Scgb1a1 in Asthma Has Been Found. Am J Respir Cell Mol Biol, 60, 615–617. 10.1165/rcmb.2019-0019ED.

39. Yogeswaran, S. & Rahman, I. 2022. Differences in Acellular Reactive Oxygen Species (ROS) Generation by E-Cigarettes Containing Synthetic Nicotine and Tobacco-Derived Nicotine. Toxics, 10. 10.3390/toxics10030134.

40. Zhu, L., An, L., Ran, D., Lizarraga, R., Bondy, C., Zhou, X., Harper, R. W., Liao, S. Y. & Chen, Y. 2019. The Club Cell Marker SCGB1A1 Downstream of FOXA2 is Reduced in Asthma. Am J Respir Cell Mol Biol, 60, 695–704. 10.1165/rcmb.2018-0199OC.

